# Single-cell analysis of ovarian myeloid cells identifies aging associated changes in macrophages and signaling dynamics

**DOI:** 10.1101/2024.06.13.598667

**Authors:** Zijing Zhang, Lu Huang, Lynae Brayboy, Michael Birrer

## Abstract

The aging of mammalian ovary is accompanied by an increase in tissue fibrosis and heightened inflammation. Myeloid cells, including macrophages, monocytes, dendritic cells, and neutrophils, play pivotal roles in shaping the ovarian tissue microenvironment and regulating inflammatory responses. However, a comprehensive understanding of the roles of these cells in the ovarian aging process is lacking. To bridge this knowledge gap, we utilized single-cell RNA sequencing (scRNAseq) and flow cytometry analysis to functionally characterize CD45^+^ CD11b^+^ myeloid cell populations in young (3 months old) and aged (14-17 months old) murine ovaries. Our dataset unveiled the presence of five ovarian macrophage subsets, including a *Cx3cr1*^low^*Cd81*^hi^ subset unique to the aged murine ovary. Most notably, our data revealed significant alterations in ANNEXIN and TGFβ signaling within aged ovarian myeloid cells, which suggest a novel mechanism contributing to the onset and progression of aging-associated inflammation and fibrosis in the ovarian tissue.

## Introduction

It has been well-established that the reproductive potential of women diminishes decades earlier than the deterioration of many of their other physiological systems. The average life expectancy for women is 79.1 years [1] while the menopause takes place at an average age of 51[2]. Furthermore, the quality of oocytes produced by women significantly decreases in their late 30s due to an increase in aneuploidy [3, 4]. This age-associated decline in female fertility has been attributed to a diminishing ovarian reserve and the alterations in the ovarian tissue microenvironment.

In both humans and mice, ovarian aging is associated with excessive deposition, as well as altered organization of collagen and hyaluronan in the ovarian stroma, signifying tissue fibrosis [5–8]. Furthermore, levels of pro- and anti-inflammatory cytokines/chemokines, including interferon-γ (IFNγ), tumor necrosis factor α (TNFα), interleukin (IL)-6, IL-10, C-C Motif Chemokine Ligand 2 (Ccl2), Ccl5, IL-4, and IL-13, have been shown to be significantly increased in aged mouse ovaries compared to that in the young ones [7, 9], indicating an increase in chronic inflammation in the tissue. These changes result, in part, in a stiff tissue texture and a dysregulated signaling environment that have been linked to impairment of follicle growth [5, 7], ovulation [10] and oocyte quality [11], as well as potentially pathological events, including the development of ovarian cancers [12, 13]. Understanding the etiology and progression of these aging-associated changes in the ovarian tissue environment is crucial for the advancement of health and reproductive longevity in women.

Immune cells are pivotal in the surveillance and regulation of the ovarian tissue microenvironment [14–16]. Myeloid cells are a group of immune cells that derive from a common myeloid progenitor in the bone marrow[17] and consist of innate immune cells, including granulocytes, monocytes, macrophages and dendritic cells[17]. They play essential roles in tissue maintenance and remodeling, as well as the orchestration of inflammatory response [17, 18]. In the mammalian ovary, macrophages are the most abundantly present myeloid cells, and are functionally indispensable to the tissue organization, vasculature development, follicle growth, and corpus luteum formation [14, 16, 19, 20].

Ovarian myeloid cell populations undergo dynamic changes over the course of aging. Studies in mice and human have reported aging-associated changes in the number and activation status of macrophages that accompany the onset of inflammation [9, 21–24], though discrepancies exist concerning the specific directions of these changes. While increased macrophage presence was observed to accompany ovarian aging in some mouse studies [9, 22], others have reported an decreased macrophage presence in aged mouse or human ovaries compared to that in the young [21, 23, 24]. Additionally, aged murine ovarian stroma also harbors a higher number of eosinophils and Ly6C^+^ monocytes compared to that in the young [8, 21]. Considering that myeloid cells are major producers of pro- and anti-inflammatory cytokines, these findings underscore the potential involvement of myeloid cells in the initiation and progression of the aging-associated inflammation.

In the past decade, advancements in single-cell RNA sequencing (scRNAseq) techniques have re-defined transcriptomic research. This powerful technique has greatly benefitted the study of ovarian aging and enabled the profiling of transcriptomic changes in aging ovarian cells, including the oocytes, granulosa cells, theca cells, and stromal cells [25, 26]. However, due to the relative scarcity of myeloid cells within the ovarian tissue, a comprehensive high-resolution analysis of their population heterogeneity and transcriptomic changes in the aging process remained largely unexplored.

In this study, we used fluorescence-assisted cell sorting (FACS) in combination with scRNAseq to obtain a focused view of myeloid cells in young (3 months old) and aged (14-17 months old) murine ovarian tissue. These cells mainly consists of macrophages, monocytes, dendritic cells (DCs), and neutrophils. Our dataset identified five distinct macrophage subsets in the murine ovarian tissue, including a Cx3cr1^low^ Cd81^hi^ subset that is almost exclusively present in the aged ovary. Our data further revealed elevated ANNEXIN and TGFβ signaling among the ovarian myeloid cells at advanced reproductive age and highlighted the potential role of neutrophils in the onset of aging-associated inflammation. Our findings provide a detailed atlas of macrophages and other myeloid cell populations in ovarian aging, laying the foundation for understanding their roles in diminished ovarian functions.

## Methods

### Animals

Virgin female C57BL/6J mice that were 2-3 months or 14-17 months of age, as well as *Cx3cr1^CreER^* and *Rosa26^floxed-tdTomato^* mice, were purchased from The Jackson Laboratory. Prior to each experiment, the estrus stage of all mice was examined using vaginal cytology. Mice at the diestrus stage were used for all experiments. Euthanasia of the animals was performed by CO_2_ inhalation. All procedures on live animals were conducted in accordance with the protocol reviewed and approved by the American Association for Laboratory Animal Science (IACUC).

### Sample processing and dissociation

Ovaries were harvested from mice aged 2-3 months old and 14-17 months. For the 2-3 months old age group, ovaries from 6 animals were assigned to 3 replicates (ovaries from 2 animals per replicate). For the 14-17 months old age group, ovaries from 5 animals were assigned to 3 replicates (2 replicates with ovaries from 2 animals and 1 replicate with ovaries from 1 animal). All samples were dissociated into a single-cell suspension in digestion buffer (1x PBS, 5% FBS, 200U collagenase IV, 200U DNaseI) using a gentleMacs^TM^ dissociator with the program 37C_m_LDK1. The suspensions were washed with 5 mL of 1x PBS with 5% FBS and 2 nM EDTA, and passed through a 100µm cell strainer to remove debris.

### Single-cell RNA sequencing and data analysis

The ovarian cell suspensions were stained with AmCyan Live/Dead stain at a 1:500 dilution in 1x PBS for 15 minutes and BioLegend TotalSeq^TM^ HTO B series hashtags at a concentration of 10 µg/mL concentration for 15 minutes. The cells were then pooled and stained with fluorochrome-conjugated primary antibodies BV650-CD45.2 (BioLegend® 109835) and PerCP/Cy5.5-CD11b (BioLegend® 101228) in 1x PBS with 5% FBS for 20 minutes. Stained cells were then washed with 1xPBS with 5% FBS and resuspended in 1x PBS with 5% FBS. FACS was performed by the Flow Core at UAMS using the BD FACSAria^TM^ III cell sorter, and single live cells with positive BV650 and PerCP/Cy5.5 staining were collected into FBS.

The encapsulation of the sorted cells and single-cell library preparation were performed by the UAMS Genomics Core using the 10x Chromium Controller and 10x Chromium NextGEM Single-Cell 3’ v3.1 kit with Featured Barcode Technology for Cell Surface Proteins following the standard protocol provided by 10x Genomics. The quality of the libraries were examined using the Agilent Fragment Analyzer capillary electrophoresis system, and the libraries were sequenced using Illumina NextSeq® 500 platform.

Quality control and alignment of the raw scRNAseq data were performed by the Genomics Core using Cell Ranger v 7.0.0, and the count matrices were generated for the young and aged libraries. The analysis of the data was performed using R 4.2.0 [27], with R package Seurat v.4 [28]. In brief, the young and aged datasets were combined, normalized, scaled, and clustered using UMAP as a reduction method [29]. Gene set enrichment analysis (GSEA) results were generated using GSEA v.4.2.1 [30] with pre-ranked gene lists produced using Seurat and R 4.2.0. Pseudotime trajectory analysis was performed using the TSCAN package v4.3 [31]. Cell-cell communication analysis was performed using CellChat package v1.6.0[32].

### Flow cytometry

The ovarian cell suspensions were stained with eBioscience^TM^ Fixable Viability Dye eFluor^TM^506 at a 1:500 dilution in 1x PBS for 15 minutes. Cells were then washed and stained with fluorochrome-conjugated primary antibodies at the recommended concentration for 20 minutes at room temperature. Monoclonal antibodies specific to mouse CD45 (BD Bioscience® 566073), CD11b (BioLegend® 101267,), F4/80 (BioLegend® 123149), CD68 (BioLegend® 137023), Ly6C (BioLegend® 128017), CX3CR1 (Biolegend® 149027), CD81 (BioLegend® 104911), MHCII (BioLegend® 107605), CD206 (Biolegend® 141731) and CD11c (BioLegend® 117365) were purchased from commercial manufacturers. The Foxp3/Transcription Factor Staining buffer set was used for Ki67 staining. Cells were resuspended in 200 µL of 1x PBS for subsequent analysis. Flow cytometry was performed by the Flow Core at UAMS using the Cytek Northern Lights^TM^ Full Spectrum Flow Cytometer. Measurements were taken on biologically independent samples. The analysis of the data was performed using SpectroFlo®, Flowjo v10.7 and Flowjo plugins UMAP v3.1. For UMAP projections of young and aged ovarian macrophages, CD45^+^F4/80^+^CD11b^+^ cells were concatenated to one FCS file before creating UMAP projections. UMAP projections were performed using default parameters.

### Fate mapping

To induce Cre recombinase activity for tracing CX3CR1+ macrophages, female *Cx3cr1^CreERT^/Rosa26^tdTomato^* mice were generated by crossing *Cx3cr1^CreER^* and *Rosa26^floxed-tdTomato^*mice. These mice were aged until they reach either 2-3 months or 14-17 months of age. To induce the recombination, tamoxifen was dissolved in corn oil and administered via oral gavage at a dose of 4 mg/day for three consecutive days. Following the last tamoxifen treatment, the mice were sacrificed at three different time points: 3 days, 10 days, and 24 days. Ovarian tissue was collected from the sacrificed mice for further analysis.

### Statistical analysis

The statistical analyses in this study were conducted using GraphPad Prism 9 software. Student’s t-test (two-sided) was employed to determine the statistical significance of the flow cytometry data. For the identification of differentially expressed genes between the two age groups, the criteria for statistical significance were defined as a minimum log fold change of 0.3 and an false discovery rate adjusted p-value lower than 0.05.

## Results

### Single-cell transcriptomic analysis determines the complex landscape of myeloid cells in young and aged murine ovaries

To examine the aging-related transcriptomic changes in ovarian myeloid cells at single-cell resolution, scRNAseq with FACS-sorted CD45^+^ CD11b^+^ cells from the ovaries of young and aged C57BL/6J virgin females was conducted. To ensure the reliability and reproducibility of our findings, 3 biological replicates for each age group were included and barcoded with hashtag oligoes (HTO).

A total of 5769 cells from the two age groups passed our quality control criteria. These cells segregated into 15 clusters based on their transcriptomic profiles (Figure 1B). The UMAP cell cluster patterns within each age group were found to be similar among the HTO-labeled replicates, indicating high robustness and reproducibility of the results (Supplemental Figure S1).

**Figure 1.**
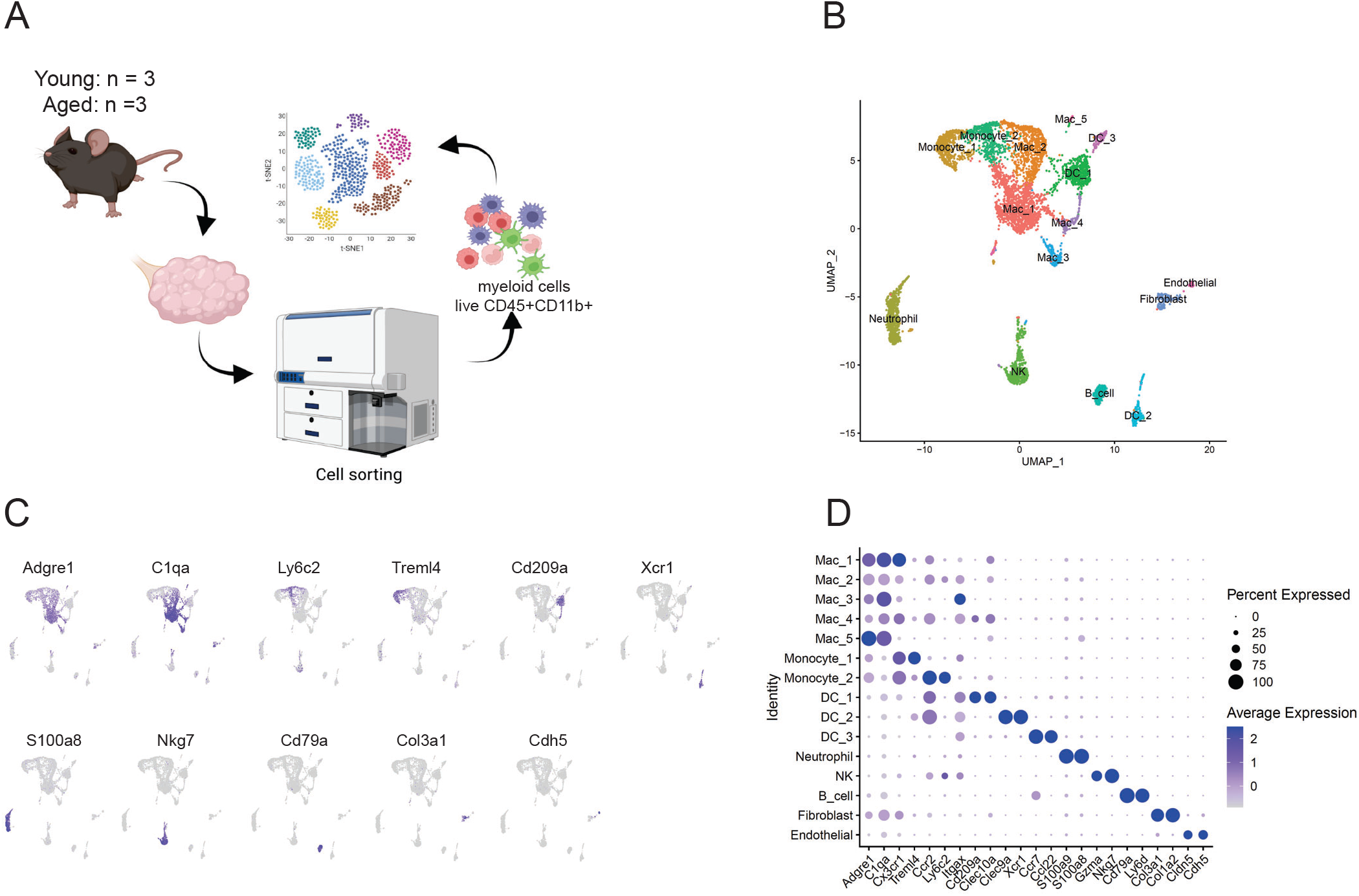
Single-cell transcriptomic analysis captured the landscape of myeloid cells in young and aged murine ovaries. **A:** Schematic drawing illustrating the scRNAseq experimental set up. **B:** The clustering of CD45+CD11b+ ovarian myeloid cells with UMAP as the dimension reduction method. The plot includes cells from all age and HTO groups. **C:** The relative expression of the representative markers of each cluster shown in the combined clustering plot. The high and low expression are represented by blue and gray colors respectively. **D:** Dotplot showing the expression of markers used for the identification of each cluster.

We annotated the clusters based on their transcriptomic signatures, and identified a total of eight cell types. The most abundant cell types include macrophages (*Adgre1*^hi^ *C1qa*^hi^), monocytes (*Adgre1*^low^ *C1qa*^low^*Cx3cr1*^hi^), dendritic cells (DC) (*Adgre1*^low^, *Cd11c*^hi^), and neutrophils (*S100a9*^hi^ *S100a8*^hi^). The dataset also captured small numbers of natural killer cells (NK) (*Gzma*^hi^, *Nk7g*^hi^), B cells (*Cd79a*, *Ly6d*), fibroblasts (*Col3a1*, *Col1a2*), and endothelial cells (*Cldn5*, *Cdh5*) (Figure 1C and D). Among these cell types, macrophages further segregated into five distinct subsets (referred to as Mac_1 to Mac_5 based on decreasing population size), monocytes consisted of two subsets distinguished by *Ly6c2* expression (*Ly6c2*^low^ Monocyte_1 and *Ly6c2*^hi^ Monocyte_2), and DCs consisted of three subsets distinguished by *Cd209a*, *Xcr1* and *Ccr7* expression (*Cd209a*^hi^ DC_1, *Xcr1*^hi^ DC_2 and *Ccr7*^hi^ DC_3) (Figure 1C and 1D).

### Ovarian macrophage population consists of multiple functionally distinct subsets

Our initial analysis focused on the ovarian macrophages since it is the most abundant among all cell types and displayed great heterogeneity in their transcriptional profiles. The dataset uncovered the presence of five macrophage subsets in the ovarian tissue (Figure 2A), each exhibits unique expression signatures indicative of different origins and functional specializations (Figure 2B).

**Figure 2.**
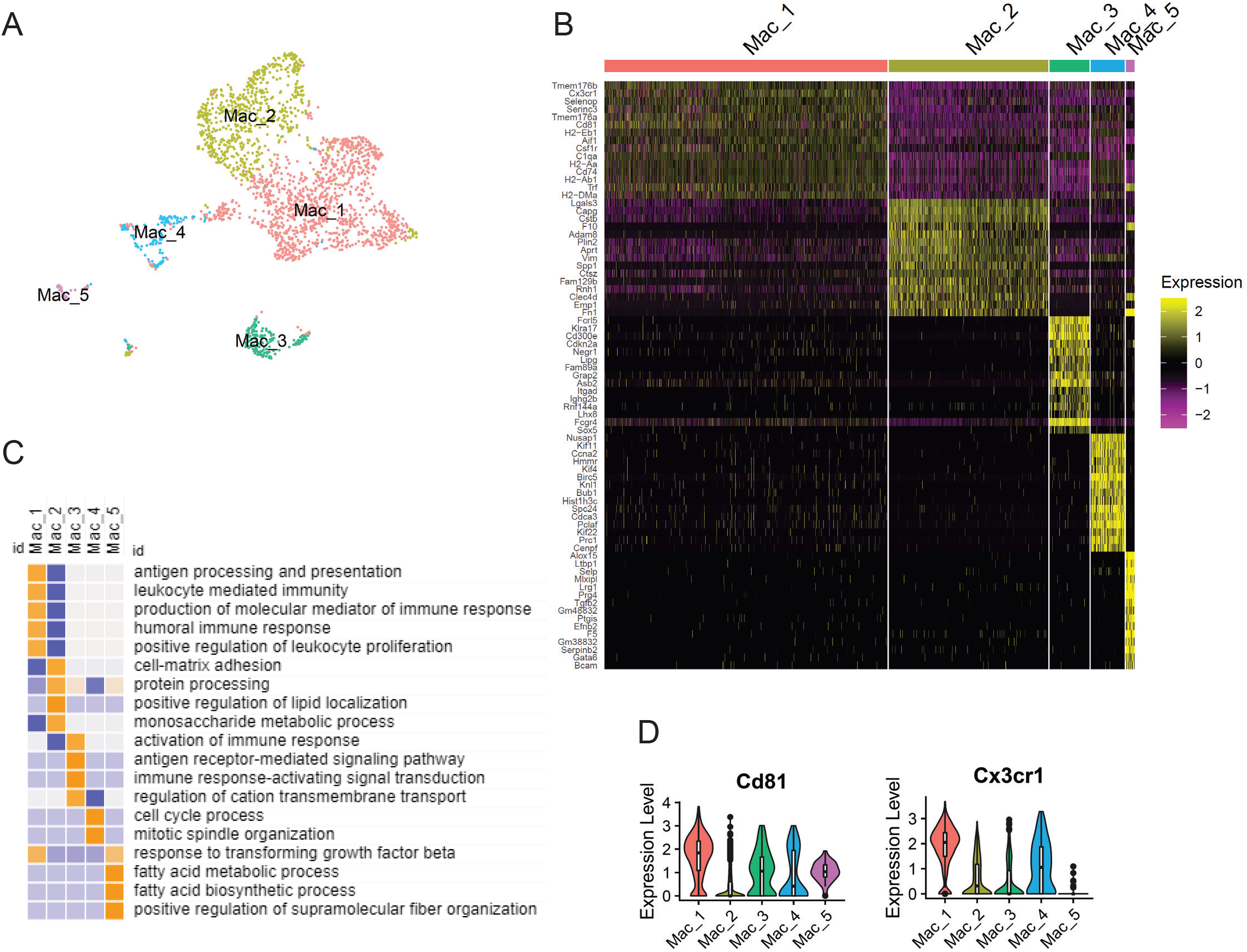
Ovarian macrophage population consists of multiple functionally distinct subsets. **A:** The clustering of ovarian macrophages that are extracted from the CD45+CD11b+ ovarian myeloid dataset. **B:** The expression of the top15 represented markers for each macrophage subpopulations ranked by fold change. **C:** The top relevant gene sets enriched in markers that are uniquely upregulated in each cluster. The color of the data point represent the Z-score of the net enrichment score (NES) of the gene set. Orange represent high value, while blue represents low value. **D:** the violin plot showing the expression of Cd81 and Cx3cr1 in the ovarian macrophage subpopulations.

Notably, Mac_5 displayed a high expression of *Gata6* and *Tgfb2* (Figure 2B), which aligned with the signature of a peritoneal macrophage population [33].

Among the four ovarian macrophage subsets (Mac_1, Mac_2, Mac_3, and Mac_4), gene sets enrichment analysis (GSEA) indicates a significant enrichment of cell cycle regulation-related genes in the transcriptomic signatures of Mac_4 (Figure 2B and 2C), indicating its identity as a cluster of proliferating macrophages. Mac_1 was highly associated with the classic macrophage function of antigen processing/presentation, while Mac_2 was associated with ECM organization. Mac_3 was associated with activation of immune response and antigen-receptor-mediated signaling, suggesting its role in regulating immune response and inflammation. The results of the gene set enrichment were validated and visualized with the module scores for the relevant gene sets (Supplemental Figure S2). Consistently, the terms “antigen presentation”, “actin and ECM organization”, “activation of immune response”, and “cell cycle” scored high in Mac_1, Mac_2, Mac_3, and Mac_4, respectively (Supplemental Figure S2).

A closer examination of the transcriptomic signatures of the macrophage subsets revealed that the most abundant three macrophage subsets, Mac_1, Mac_2 and Mac_3, could be distinguished based on their expression of the chemokine receptor *Cx3cr1* and the tetraspanin *Cd81* (Figure 2D). Mac_1 exhibited high expression of both *Cx3cr1* and *Cd81* (Cx3cr1^hi^, Cd81^hi^), Mac_2 displayed low expression of both markers (*Cx3cr1*^low^, *Cd81*^low^), and Mac_3 showed low *Cx3cr1* and high *Cd81* expression (*Cx3cr1*^low^, *Cd81*^hi^) (Figure 2C).

Since the functions of macrophages are heavily influenced by their activation status, we examined the expression pattern of markers that are commonly associated with classical (M1) and alternative (M2) macrophage activation status among the ovarian macrophage subsets [34]. Interestingly, a mixture of M1 and M2-associated markers was found to be expressed in all macrophage subsets (Supplemental Figure S3). Thus, none of the ovarian macrophage subpopulations show obvious bias towards M1 or M2 polarization.

### Cx3cr1^hi^Cd81^hi^ Mac_1 represents a self-maintained low turnover ovarian macrophage subset

To validate the scRNAseq findings, we performed flow cytometry analysis with CD68^+^ F4/80^+^ ovarian macrophages (Supplemental Figure S4). Consistent with the sequencing data, ovarian macrophages segregate into distinct CX3CR1^hi^CD81^hi^, CX3CR1^low^CD81^low^ and CX3CR1^low^CD81^hi^ subsets that correspond to Mac_1, Mac_2 and Mac_3 (Figure 3A). Additionally, the analysis captured a CX3CR1^hi^ CD81^low^ macrophage subset not observed in the scRNAseq data (Figure 3A), which may represent a transitional population.

**Figure 3.**
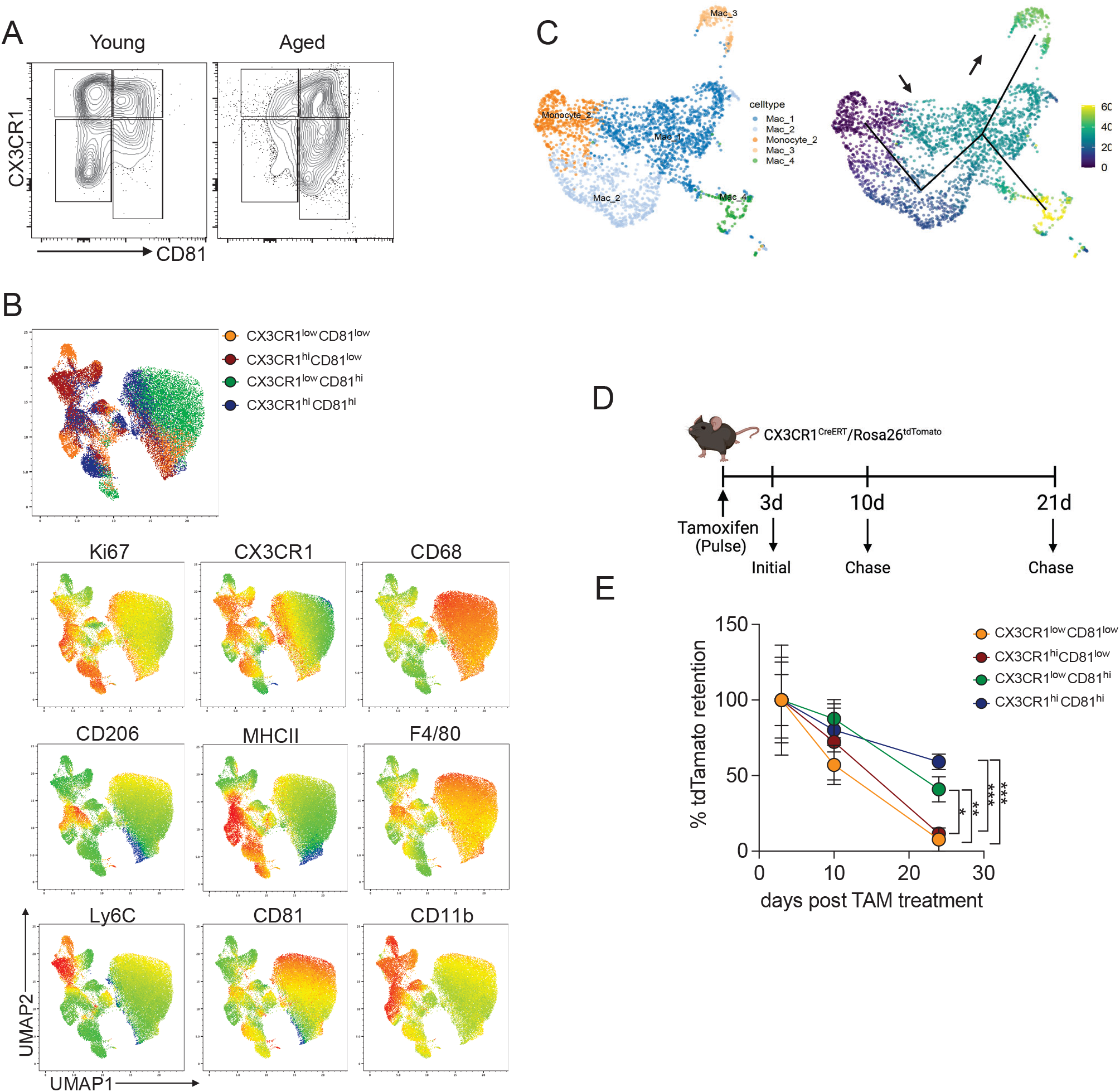
CX3CR1^hi^CD81^hi^ macrophages represents a stable self-maintaining ovarian macrophage subset. **A:** Representative flow cytometry plot of CX3CR1 and CD81 expression in ovarian macrophages in young and aged murine ovary. **B:** Clustering of pooled CD68^+^ F4/80^+^ according to the expression of Ki67, CX3CR1, CD68, CD206, F4/80, MHCII, CD81, and Ly6C. UMAP was used as the dimension reduction method. The individual plots showed the four CX3CR1/CD81 defined macrophage subsets, and the expression of Ki67, CX3CR1, CD68, CD206, F4/80, MHCII, CD81, and Ly6C respectively. **C:** Pseudotime trajectory model calculated based on the clustering of Monocyte_2, Mac_1, Mac_2, Mac_3 and Mac_4. **D:** Schematic drawing illustrating the fate mapping experiment with CX3CR1^CreERT^/Rosa26^tdTomato^ mice. **E**: Percentage of total tdTomato positive cells in the CX3CR1/CD81 defined macrophage subsets 3 days (n = 3), 10 days (n = 7) and 24 days (n = 6) post TAM treatment.

The characterization of these ovarian macrophage subsets using dimensional flow cytometry analysis led to several interesting observations. First, CX3CR1^hi^CD81^low^ and CX3CR1^low^CD81^low^ (Mac_2) subsets exhibited high expression of Ly6C (Figuire 3B) and lower levels of F4/80 and CD68, which is consistent with macrophages that are newly differentiated from monocytes [35]. Secondly, CX3CR1^low^CD81^hi^ (Mac_3) cells and CX3CR1^hi^CD81^hi^ (Mac_1) cells showed high expression of F4/80 and CD206 (Figure 3B), suggesting their identities as tissue-resident macrophages [35]. Thirdly, although Ki67 expression was observed in all macrophage subsets, the highest Ki67 expression mostly overlap with CX3CR1^hi^CD81^hi^ (Mac_1) (Figure 3B), which suggests Mac_1 as a self-maintained macrophage population with a high number of proliferative cells [36]. The expression patterns of markers observed in the flow cytometry analysis largely aligned with those observed in the scRNAseq dataset (Supplemental Figure S5). Taken together, our data indicated that Mac_1 represents a mature, self-maintained ovarian macrophage subset, while Mac_2 and CX3CR1^hi^CD81^low^ macrophages appeared to have more recently differentiated from monocytes.

To verify the relationships between the ovarian macrophage subsets, we generated a pseudotime trajectory incorporating the four ovarian macrophage subsets and the *Ly6c2*+ monocyte subset (Monocyte_2) using the TSCAN package [31]. Monocyte_2 was set as the root of the trajectory, as it represents the least differentiated state. The pseudotime model identified Mac_3 and Mac_4 as two end point clusters, inferred as derivatives of the Mac_1 subset (Figure 3C). Mac_2 was identified as an intermediate state between Monocyte_2 and Mac_1 (Figure 3C). Overall, the pseudotime analysis provided further evidence supporting the close association of Mac_1 with Mac_3 and the proliferative macrophage population (Mac_4).

To validate the self-maintaining status of Mac_1, we conducted fate-mapping analysis of the CX3CR1/CD81-defined ovarian macrophage subsets. All ovarian macrophages express CX3CR1 and thus can be labeled in the Cx3cr1^CreERT^ inducible fate mapping mouse model, where a tamoxifen-inducible Cre recombinase is expressed under the control of the endogenous Cx3cr1 promoter (Figure 3D). To track all ovarian macrophage subsets and measure their turnover rate in aged mice, the *Cx3cr1*^CreERT^ mouse was crossed to a *Rosa26* ^tdTomato^ reporter mouse and a pulse-chase experiment was conducted. For the four macrophage subsets, we calculated the ratio of % tdTomato+ cells at the later time points to % tdTomato+ cells at day 3 post tamoxifen treatment. Thus, a reduced % tdTomato retention indicates a decrease in tdTomato+ cells over time. Notably, while all four macrophage subsets were labeled by tdTomato, the % tdTomato retention decreased dramatically in both the CX3CR1^low^CD81^low^ (Mac_2) and the CX3CR1^hi^CD81^low^ subsets. However, the CX3CR1^low^CD81^hi^ (Mac_3) and CX3CR1^hi^CD81^hi^ (Mac_1) subsets displayed much slower decreases in % tdTomato retention (Figure 3E). These data indicate that the four ovarian macrophage subsets exhibited differences in turnover properties, with Mac_1 and Mac_3 subsets being relatively long-lived in the aged ovary, supporting the stable, self-maintaining property of Mac_1 and the relationship between Mac_1 and Mac_3.

### The CX3CR1^low^ CD81^hi^ Mac_3 macrophage subset drastically expands in aged murine ovaries

The comparison of scRNAseq data between the two age groups unveils that, while the presence of Mac_1, Mac_2, Mac_4 and Mac_5 did not show statistically significant differences across the two age groups, the Mac_3 subset was almost exclusively present in the reproductively old tissue (Figure 4A). This observation was confirmed by the flow cytometry data, as the proportion of CX3CR1^low^CD81^hi^ subset in total macrophage population increased significantly in aged ovarian tissue (Figure 3A and 4B).

**Figure 4.**
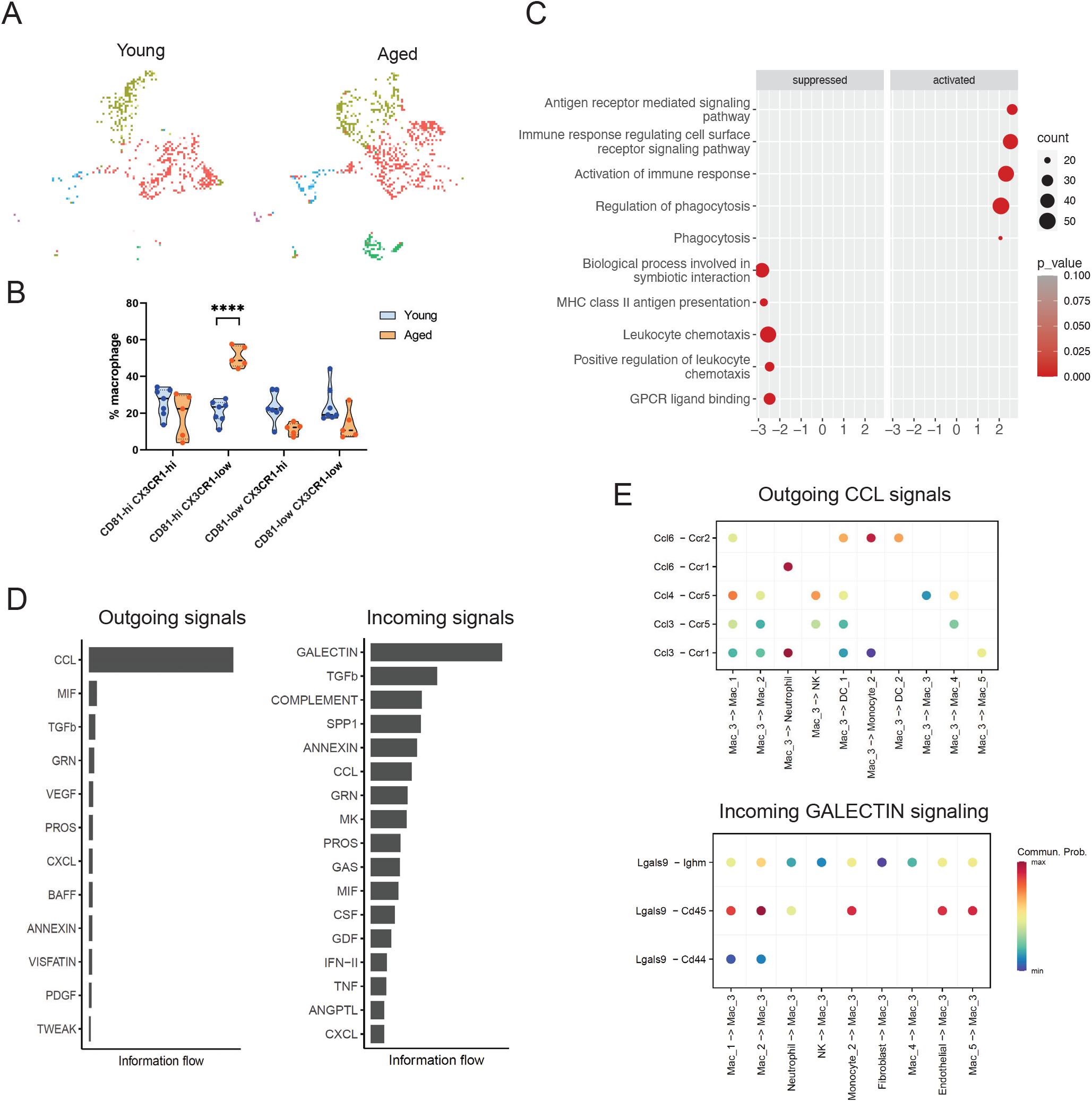
A CX3CR1^low^ CD81^hi^ macrophage subset is uniquely present in aged murine ovaries. **A:** The clustering of ovarian macrophages split by age, shown with UMAP as the reduction method. **B:** The quantification of the CX3CR1^hi^CD81^low^, CX3CR1^hi^CD81^hi^, CX3CR1^low^CD81^hi^, CX3CR1^low^CD81^low^ macrophage subpopulations in young (n = 5) and aged (n = 7) murine ovaries based on the flow cytometry analysis. For CX3CR1^low^CD81^hi^ macrophages, p = 0.00006 (aged vs young). **C:** The top relevant terms in the GSEA results of the expression signatures of the Mac_3 macrophage subset compared to Mac_1, Mac_2, Mac_4 and Mac_5. **D:** The top pathways in the outgoing and incoming signaling from and to the Mac_3 subset. The ranking was performed based on information flow. Higher value indicates higher overall signaling strength. **E:** Bubble plot showing the relative strength of specific ligand-receptor interaction within the outgoing CCL and incoming GALECTIN pathways from and to the Mac_3 subset. The color of the bubble represent the relative communication probability.

To gain insights into the properties of the Mac_3 subset, we analyzed its transcriptomic profile using GSEA[30]. This analysis revealed significant downregulation of genes related to chemotaxis, and a simultaneous upregulation of genes associated with antigen receptor signaling, immune response activation and phagocytosis (Figure 4C). These results suggest Mac_3 as a subset of low-mobility macrophages with increased phagocytic activities. Remarkably, Mac_3 exhibits high expression of *Pparg* (Supplemental Figure S3), a transcription regulator essential to promote lipid metabolism in macrophages [37]. This suggests that Mac_3 may engage in a metabolic program that enables efficient lipid utilization.

To explore the influence of Mac_3 within the tissue microenvironment, we inspected its interactions with other cell clusters within the dataset. Employing the CellChat R package, we inferred the signaling activities to and from Mac_3 in the aged myeloid cell populations based on ligand-receptor interactions [32]. Intriguingly, the analysis reveals that the outgoing signals from Mac_3 are predominantly that of the CCL pathway, while TGFβ, vascular endothelial growth factor (VEGF), and CXCL are also among the top outgoing signaling pathways with relatively high information flow. Conversely, Mac_3 mainly receives signaling through the GALECTIN, TGFβ, and COMPLEMENT pathways (Figure 4D). A closer examination of the CCL signaling from Mac_3 indicates that the subset mainly produces *Ccl6*, *Ccl3* and *Ccl4*. And the strongest communication takes place between Mac_3 and the neutrophils through *Ccl6*-*Ccr1* and *Ccl3*-*Ccr1* interactions (Figure 4E). GALECTIN signaling received by Mac_3 predominantly originates from other macrophage subsets and Monocyte_2, featured with *Lgals9*-CD45 interactions (Figure 4E).

### Aged ovarian myeloid cells are featured with intensified ANNEXIN and TGFβ signaling

To explore the temporal changes in the ovarian myeloid cells and elucidate their roles in the ovarian aging process, we analyzed and compared the cell-cell communication network in the two age groups based on the single-cell transcriptomic data. Overall, we observed an increase in both the number and the strength of cell-cell interactions among myeloid cells in the aged ovary compared to that in the young, with an exception being the outgoing signaling activities from Mac_2 (Figure 5A).

**Figure 5.**
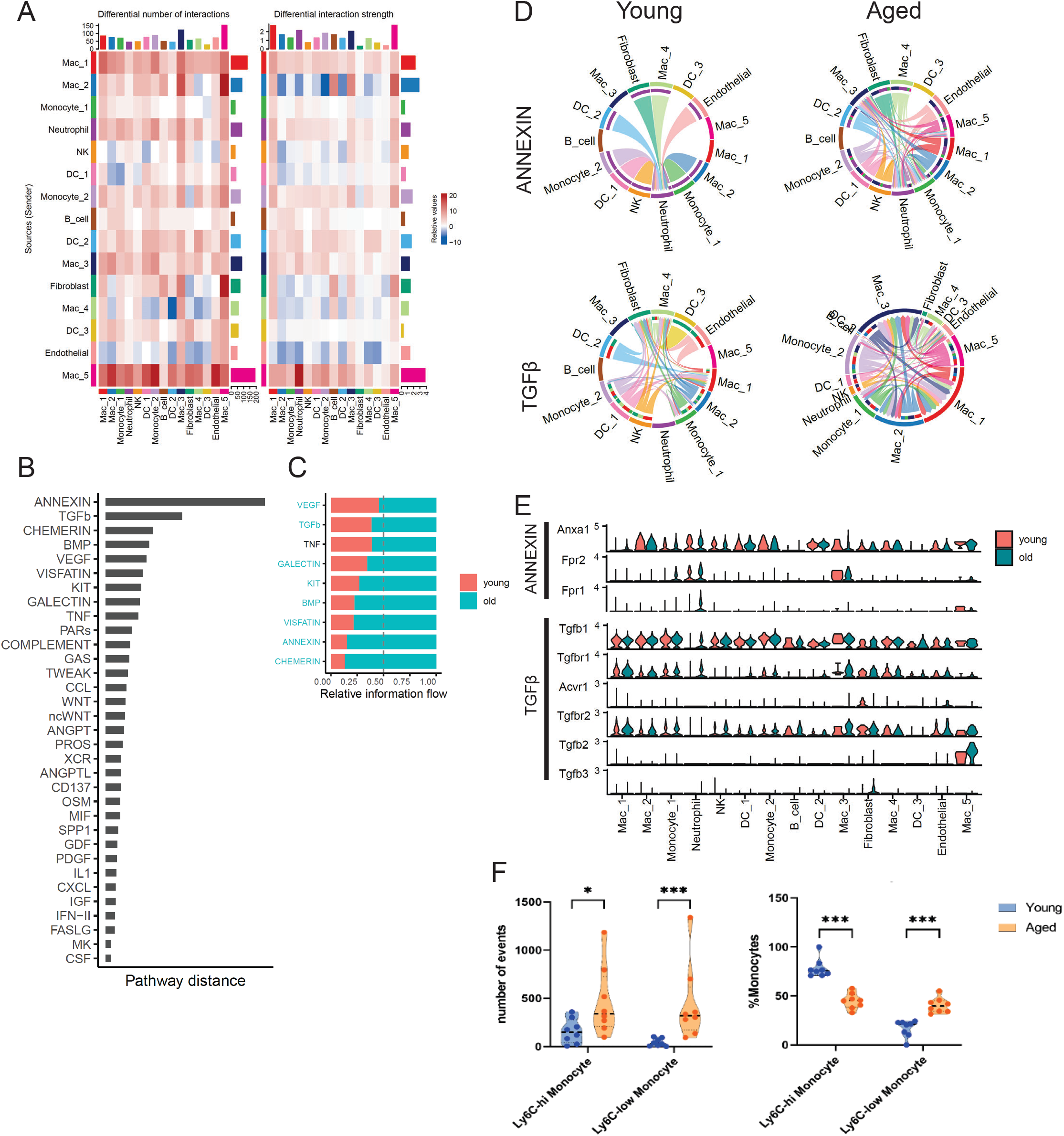
Elevated ANNEXIN and TGFβ signaling are observed among aged ovarian myeloid cells. **A:** The relative changes of the number and strength of signaling communication between different cell clusters in aged vs young myeloid cells. The color of the cells represent the direction of changes. Red represents increase while blue represent decrease. The bars at the end of columns and rows represents the column or row sums respectively. **B:** The ranking of pathways that are present in the communication network among both the young and aged myeloid cells based on is functional distinction between the two age groups. Higher value indicate greater changes in strength, sender/receiver identification, and dominant ligand-receptor pairs. **C:** Relative information flow of the top ranked pathways in panel B in young and aged communication network. The length of the bar represents relative signaling strength. **D:** The chord plot that visualize the sender and receivers of ANNEXIN and TGFβ signaling in young and aged communication networks. The color of the outer and inner bars represents the identity of the sender and receiver of the signaling respectively. The size of the bars represents the signal strength. **E:** The violin plot that shows the expression of ligand and receptors relevant to the ANNEXIN and TGFβ pathways in all cell clusters split by age. **F:** The quantification of the count and proportions of Ly6C^hi^ and Ly6C^low^ monocytes in young (n = 8) and aged (n = 8) murine ovary based on the flow cytometry analysis. Adjusted p-values for the comparisons from left to right are: Ly6C-hi monocyte absolute count aged vs young, p = 0.028127; Ly6C-low monocyte absolute count aged vs young, p = 0.000466; Ly6C-hi monocyte as %Monocyte aged vs young, p = 0.000155; Ly6C-low monocyte as %Monocyte aged vs young, p = 0.000155.

Pathways that are present in the signaling network of both age groups may exhibit significant functional differences due to alterations in the senders and receivers of the signaling and the predominant ligand-receptor pairs[32]. We ranked pathways shared between the two age groups based on their functional deviations in the young and aged cell-cell communication networks. Strikingly, the most prominent age-associated alterations were observed in ANNEXIN and TGFβ pathways (Figure 5B). Additionally, BMP, VEGF, VISFATIN, TNF pathways also ranked among the most functionally distinct between the two age groups (Figure 5B). Interestingly, all these markedly altered pathways exhibited significantly intensified signaling activities in the older age group (Figure 5C).

We delineated the most significantly altered ANNEXIN and TGFβ signaling pathways within young and aged cells, and found drastic changes in both the sources and targets of these signaling interactions (Figure 5D). Among young ovarian myeloid cells, ANNEXIN signaling originates from multiple sources, including Mac_2, Mac_4, DC_1, DC_2, monocytes, and neutrophils, with neutrophils being the sole target. In aged ovarian myeloid cells, however, Mac_1 and Mac_5 emerged as additional sources of ANNEXIN signaling, while the target cell types expanded to include neutrophils, Mac_3, Mac_5 and Monocyte_1 (Figure 5D). Regarding the TGFβ pathway, there was a marked increase in overall signaling activities and a drastic expansion of incoming signaling to macrophage subsets including Mac_1, Mac_2 and Mac_3 (Figure 5D).

Upon Examination of the relevant ligand and receptor expression, it became evident that the changes in ANNEXIN signaling activities were mainly a result of increased expression of Anxa1 in aged Mac_1 and Mac_5, and upregulation of *Anxa1* receptor *Fpr2* in Monocyte_1, Mac_3 and Mac_5 (Figure 5E). Changes in TGFβ signaling can be attributed to a general elevation in *Tgfb1* expression across macrophage subsets, along with increased expression of *Tgfbr1* and *Tgfbr2* within the macrophages (Figure 5E).

Notably, Anxa1 encodes Annexin A1 (ANXA1), an anti-inflammatory protein that orchestrate the resolution of inflammation. ANXA1 is a chemoattractant of monocytes[38–40]. Flow cytometry analysis showed that the aging-associated increase in ANNEXIN signaling to Monocyte_1 is accompanied by significant increase in both the number and proportion of Monocyte_1 in aged ovary (Supplemental Figure S4, Figure 5F), providing further evidence for the elevated signaling activities.

## Discussion

In this study, five distinct ovarian macrophage subpopulations have been identified through single-cell transcriptomic analysis. Based on the expression of Ly6C, CD81 and CD11b, as well as their turnover rates in the aging ovarian tissue, the CX3CR1^hi^ CD81^hi^ Mac_1 and CX3CR1^low^ CD81^low^ Mac_2 subsets may correspond to the long-lived F4/80^hi^ CD11b^int^ and short-lived F4/80^int^ CD11b^hi^ ovarian macrophage subtypes identified by Li et al using flow cytometry in a recent publication[41]. Mac_1 was characterized as a mature, self-maintaining macrophage subset that resembles tissue-residential macrophages.

Interestingly, our data revealed a unique subset of CX3CR1^low^CD81^hi^ macrophages in aged ovarian tissue. The high expression of Pparg in this subset indicates a distinct adaptation to increased lipid metabolism. Aging and obesity often result in a similar inflammatory milieu in the ovarian tissue, where aging is associated with increased visceral adipose accumulation and altered lipid profiles in and around the ovary [42, 43]. It is thus plausible that the upregulation of Pparg expression in this macrophage subset signifies an adaptation to the ovarian tissue environment at advanced reproductive age. Moreover, it is known that PPARγ activation can enhance Fc-receptor-mediated phagocytosis in macrophages [44–47]. Aligning with this, genes associated with the regulation of phagocytosis are found to be upregulated in this specific macrophage subpopulation, suggesting increased phagocytic activities. Another intriguing discovery about the CX3CR1^low^CD81^hi^ macrophage subset is the predominance of CCL pathway in its signaling output. This subset expresses high level of *Ccl3* and *Ccl4*, known as macrophage inflammatory proteins (MIPs) that can stimulate the production of pro-inflammatory cytokines, including TNFα, IL-1β, and IL-6, within macrophages[48]. Additionally, it expresses *Ccl6*, a chemoattractant for monocytes and macrophages[49, 50] that has been linked to pulmonary inflammation and fibrosis[51, 52]. These findings suggest a plausible role for the CX3CR1^low^CD81^hi^ macrophage subset in the development of age-related chronic inflammation and fibrosis within ovarian tissue. It is conceivable that this macrophage subset might be associated with the F4/80+ group of multi-nuclei giant cells identified through histological studies in aged ovarian tissue [7, 53]. However, further investigations will be imperative to ascertain the relationship between these two subsets.

In addition to the emergence of the CX3CR1^low^CD81^hi^ macrophage subset, one of the most conspicuous changes associated with aging in ovarian tissue is the profound alterations observed in the cell-cell communication networks, particularly within the ANNEXIN and TGFβ pathways. Studies into the mechanisms and progression of inflammation suggested a potential interconnection between the upregulation of these two pathways. ANXA1, a pivotal player in the resolution of inflammation, performs crucial roles during infection and wound healing. It is released by infiltrating myeloid cells, particularly neutrophils and macrophages[54], to attenuate further neutrophil infiltration, recruit monocytes, and activate the expression of anti-inflammatory cytokines and TGFβ1 in macrophages [54, 55]. Our data aligned with this concept, revealing an age-related increase in the presence of Monocyte_1 (Figure 5F), and a high expression of *Tgfb1* in Mac_3 (Figure 5E), both of which are targets of the heightened ANNEXIN signaling in aged ovarian tissue. Given that TGFβ signaling promotes the alternative activation of macrophages, the increased ANNEXIN signaling may contribute to the rise of M2-like macrophages in the reproductively aged ovary [9, 21]. Consequently, our findings cast ANNEXIN signaling as a compelling avenue warranting further investigation and offering novel therapeutic potentials.

Furthermore, neutrophils emerge as a major source of ANXA1 during wound healing [54]. Our single-cell data not only reflects an increase in the presence of neutrophil in aged ovarian tissue (Supplemental Figure S6A), but also indicates an elevation in the intensity of both incoming and outgoing ANNEXIN signaling within aged neutrophils (Supplemental Figure S6B). In each ovarian cycle, the release of the egg leads to wounding of the ovarian surface and subsequent wound healing that recruit neutrophils. The inability to effectively clear neutrophils during this process could potentially lead to the accumulation of these granulocytes and their detrimental contents in the ovarian tissue over time, which may contribute to the onset of chronic tissue inflammation and fibrosis. Our data underscores the role of neutrophils as a previously overlooked cellular cohort that might be intricately linked to ovarian aging.

Finally, our scRNAseq dataset also captured a small GATA6^+^ macrophage subset resembling the peritoneal macrophages. Intriguingly, GATA6^+^ peritoneal macrophages have been shown to be recruited for tissue repair in other organs within the peritoneal cavity, such as the liver and intestine [56, 57]. Considering that the mammalian ovary undergoes tissue damage and repair in each ovarian cycle, it is possible that the peritoneal macrophages are recruited and play a functional role in the ovarian tissue repair process. Further investigation into the recruitment and functional importance of peritoneal macrophages in ovarian tissue repair would be an interesting area of research.

In addition to aging, immunological changes in ovarian tissue have been increasingly associated with various pathological conditions, such as diminished ovarian reserve [22], ovarian cancers [13], and polycystic ovary syndrome [58]. The scRNAseq dataset generated in this study has the potential to provide valuable insights into the understanding of these diseases as well. The findings of this study may have significant translational value if they can be validated in ovarian specimens from human patients.

## Acknowledgement

This study is generally funded by the Global Consortium of Reproductive Longevity and Equality through Buck Institute to ZZ, as well as the Start-up package from UAMS to MB. LH is supported by National Institutes of Health (NIH)/National Institute of General Medical Sciences grants (P20GM103625 and P30GM145393 to through the Center for Microbial Pathogenesis and Host Inflammatory Responses at UAMS), American Lung Association (CA-828143), Arkansas Bioscience Institute, Sturgis Foundation and Start-up package from College of Medicine at UAMS.

## Authors’ contribution

ZZ is responsible for the design and execution of the experiments, data analysis, and the writing of the manuscript. LB advised the design of the experiments and writing of the manuscript. LH and MB supervised the study, and advised the design/execution of the experiments and the writing of the manuscript.

## Notes

### Competing Interest Statement

The authors have declared no competing interest.

