## Supplemental Figures for "Single-cell analysis of ovarian myeloid cells identifies aging associated changes in macrophages and signaling dynamics"

Supplementary Figure 1

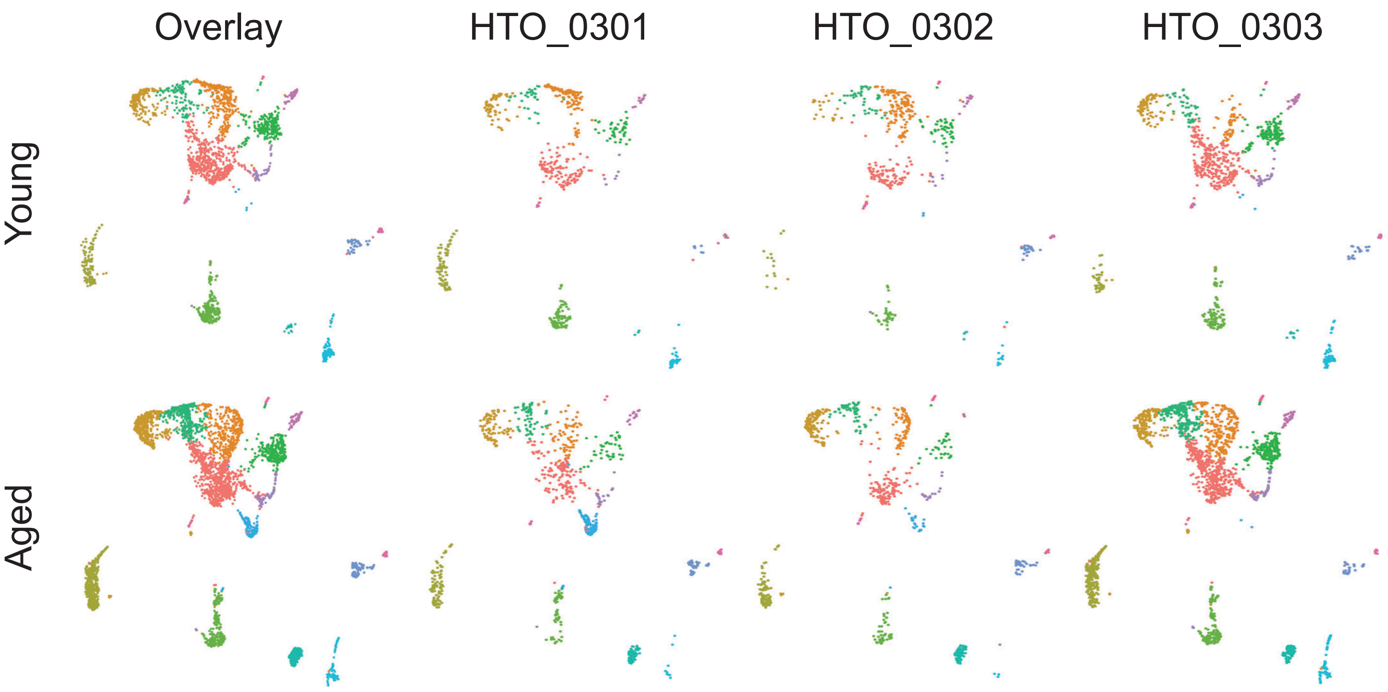

Supplementary Figure 2

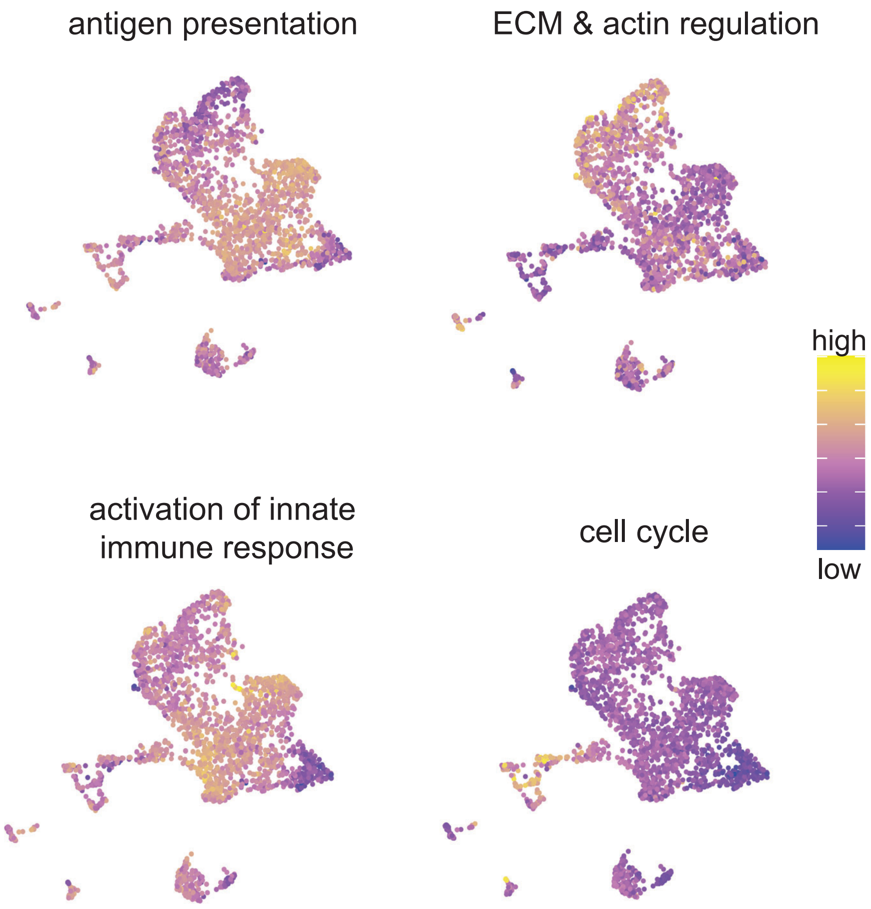

### Supplementary Figure 3

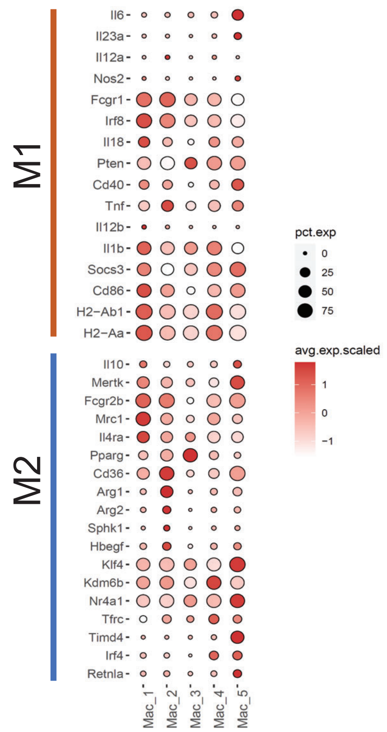

Supplementary Figure 4

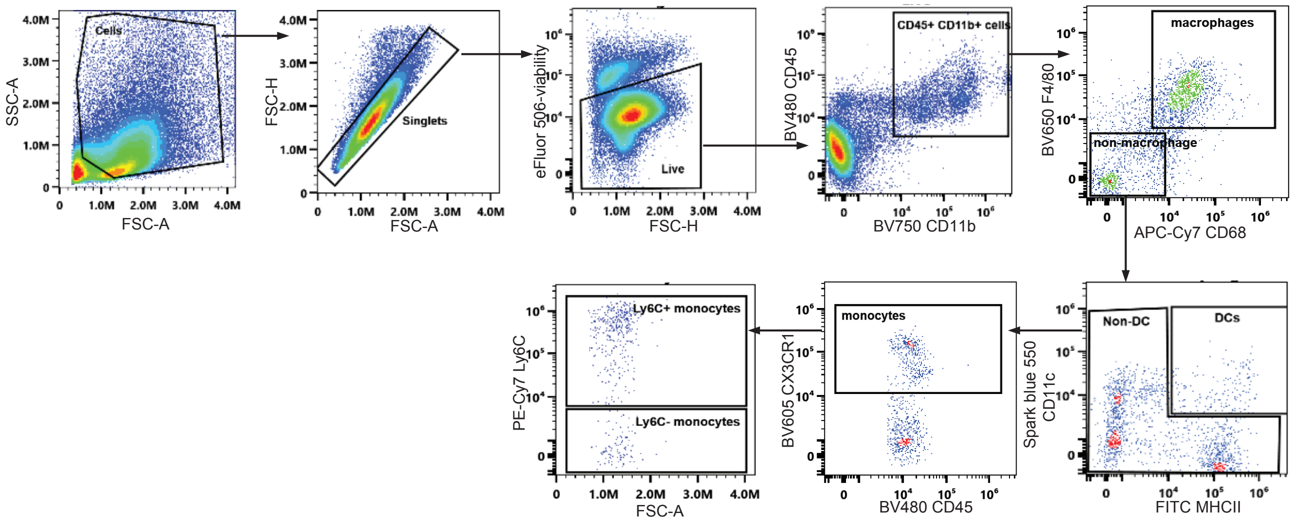

Supplementary Figure 5

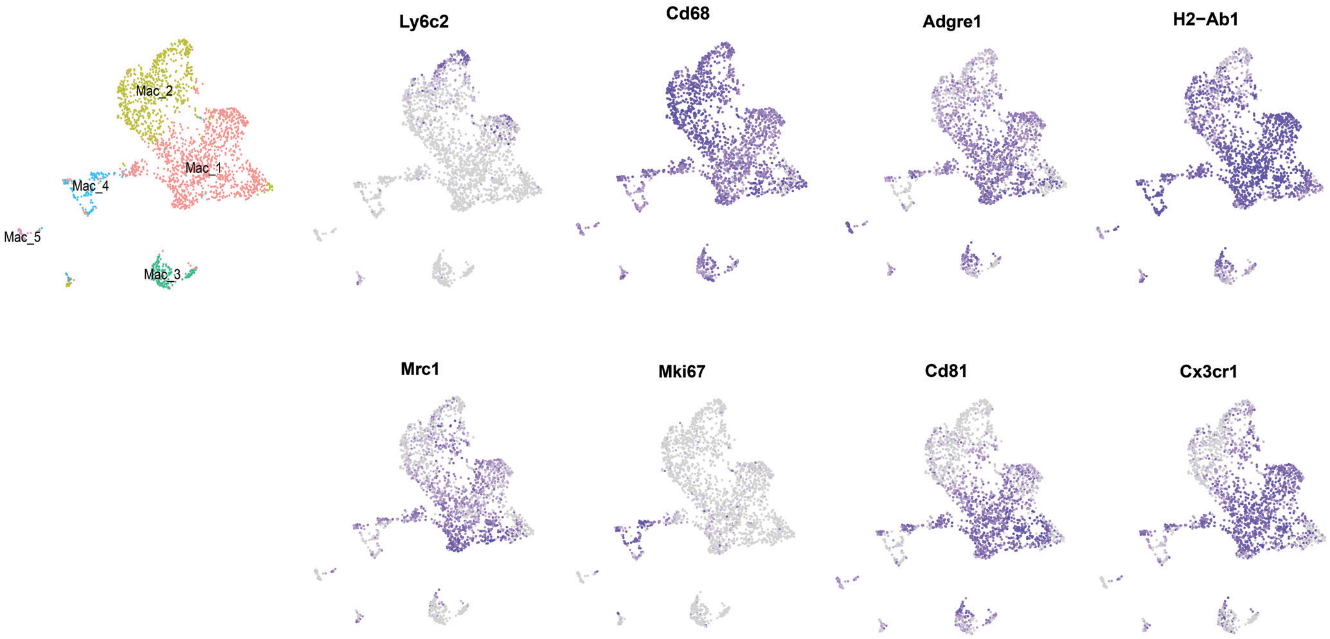

### Supplementary Figure 6

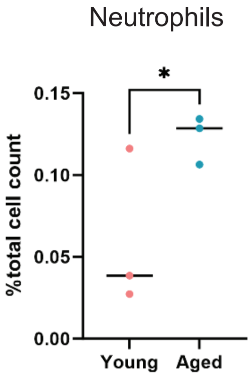


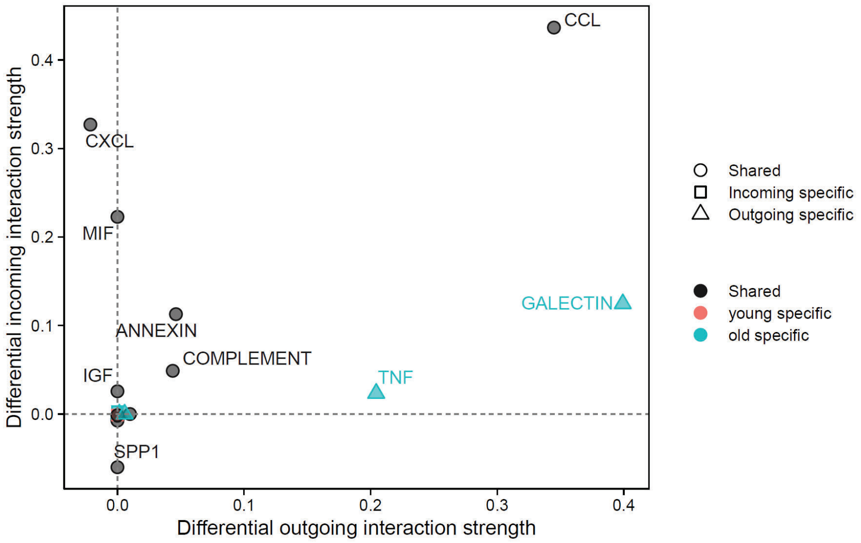
