## Supplemental Figure Legends for "Single-cell analysis of ovarian myeloid cells identifies aging associated changes in macrophages and signaling dynamics"

### Supplemental Figures

**Supplemental Figure S1.** The clustering of CD45+CD11b+ ovarian myeloid cells in each age and HTO label groups.

**Supplemental Figure S2.** The dot plots of relative gene set expression levels calculated as metric scores in ovarian macrophage subpopulations. The high and low scores are represented by yellow and blue color respectively. The metric scores are calculated as described in Tirosh et al 2016[1].

**Supplemental figure S3**. The expression of markers commonly associated with M1-like and M2-like macrophages. The size of the dots represent the percent of cells in a cluster that express the marker. The shade of the dots represent the scaled average expression of the marker in a cluster.

**Supplemental Figure S4.** The gating strategy for macrophages and monocytes in flow cytometry analysis.

**Supplemental Figure S5.** The expression of Ly6c2, Cd68, Adgre1 (F4/80), H2-Ab1 (MHC-II), Mrc1 (CD206), Mki67, Cd81, Cx3cr1 in ovarian macrophage subsets as reflected by the scRNAseq data. Blue indicates high expression.

**Supplemental Figure S6. The neutrophil population increase in number and send out more ANNEXIN signaling in aged ovarian tissue. A:** The quantification of neutrophil cell numbers in the different HTO labeled biological replicates based on the scRNAseq dataset. **B:** The relative changes in incoming and outgoing signaling strength of different pathways to and from neutrophils in old vs young communication network. The colors of the points indicate whether the pathway exist specifically in the young or the old, or in both young and old communication network. The shape of the points represent the incoming/outgoing specificity of the pathway.
